# Neural evidence for the prediction of animacy features during language comprehension: Evidence from MEG and EEG Representational Similarity Analysis

**DOI:** 10.1101/709394

**Authors:** Lin Wang, Edward Wlotko, Edward Alexander, Lotte Schoot, Minjae Kim, Lena Warnke, Gina R. Kuperberg

## Abstract

It has been proposed that people can generate probabilistic predictions at multiple levels of representation during language comprehension. We used Magnetoencephalography (MEG) and Electroencephalography (EEG), in combination with Representational Similarity Analysis (RSA), to seek neural evidence for the prediction of animacy features. In two studies, MEG and EEG activity was measured as human participants (both sexes) read three-sentence scenarios. Verbs in the final sentences constrained for either animate or inanimate semantic features of upcoming nouns, and the broader discourse context constrained for either a specific noun or for multiple nouns belonging to the same animacy category. We quantified the similarity between spatial patterns of brain activity following the verbs until just before the presentation of the nouns. The MEG and EEG datasets revealed converging evidence that the similarity between spatial patterns of neural activity following animate constraining verbs was greater than following inanimate constraining verbs. This effect could not be explained by lexical-semantic processing of the verbs themselves. We therefore suggest that it reflected the inherent difference in the semantic similarity structure of the predicted animate and inanimate nouns. Moreover, the effect was present regardless of whether a specific word could be predicted, providing strong evidence for the prediction of coarse-grained semantic features that goes beyond the prediction of individual words.

**Significance statement:** Language inputs unfold very quickly during real-time communication. By predicting ahead we can give our brains a “head-start”, so that language comprehension is faster and more efficient. While most contexts do not constrain strongly for a specific word, they do allow us to predict some upcoming information. For example, following the context, “they cautioned the…”, we can predict that the next word will be animate rather than inanimate (we can caution a person, but not an object). Here we used EEG and MEG techniques to show that the brain is able to use these contextual constraints to predict the animacy of upcoming words during sentence comprehension, and that these predictions are associated with specific spatial patterns of neural activity.

## Introduction

Probabilistic prediction is proposed to be a fundamental computational principle underlying language comprehension (Kuperberg and Jager, 2016). Evidence for this hypothesis comes from the detection of anticipatory neural activity prior to the appearance of strongly predicted incoming words (e.g. Wicha et al., 2004; Wang et al., 2018). In natural language, however, contexts that predict specific words appear relatively infrequently (Luke and Christianson, 2016). Therefore, for prediction to play a major role in language processing, comprehenders must be able to use contextual constraints to predict features that characterize multiple upcoming inputs. Here, we ask whether comprehenders can use the constraints of verbs to predict semantic features associated with the *animacy* of upcoming nouns during discourse comprehension.

The ability to distinguish between animate and inanimate entities is fundamental to human cognition (Caramazza and Shelton, 1998; Nairne et al., 2017) and to the structure of language (Dahl, 2008). Verbs can constrain for the animacy of their arguments (McCawley, 1968; Jackendoff, 1993), and these constraints can lead to anticipatory behavior during online language comprehension (Altmann and Kamide, 1999). Moreover, a larger event-related potential (ERP) response (the N400) is evoked by nouns that mismatch (versus match) these animacy constraints (Paczynski and Kuperberg, 2011, 2012; Szewczyk and Schriefers, 2011), and neural effects to mismatching inputs can be detected even before the animacy features of upcoming arguments become available (Szewczyk and Schriefers, 2013). Here, we sought direct neural evidence for the prediction of animacy features in the absence of any bottom-up input by exploiting the inherent difference in the *semantic similarity structure* of animate and inanimate nouns.

Animate entities share more co-occurring semantic features, which are more strongly intercorrelated, than inanimate entities (McRae et al., 1997; Zannino et al., 2006). For example, the animate words, “swimmer” and “pilot”, share more co-occurring semantic features (e.g. <can move>, <can breathe>, <sentient>) than the inanimate words, “paper” and “water”, which have more distinct features (e.g. <thin> for “paper”, <drinkable> for “water”). Modeling work using both connectionist (Rogers and McClelland, 2008) and Bayesian (Kemp and Tenenbaum, 2008) frameworks shows that patterns of covariation amongst internal representations of concepts can account for the emergence of categorical taxonomic structure. Therefore, these differences in the semantic similarity of animate and inanimate entities can also explain why the overall category, <Inanimate>, subsumes a larger number of subordinate semantic categories (e.g. <vegetables>, <furniture>, <tools>) than the overall category, <Animate> (Garrard et al., 2001).

In the brain, semantic features are thought to be represented within widely distributed networks (Martin, 2016; Huth et al., 2016). Thus, differences between *animate* and *inanimate* concepts in their internal semantic similarity structures can give rise to differences in similarity amongst the spatial patterns of neural activity associated with their processing. These differences can explain specific patterns of category-specific deficits in patients with non-focal neuropathologies (Devlin et al., 1998; Tyler and Moss, 2001). They can also be detected using Representational Similarity Analysis (RSA) (Kriegeskorte et al., 2008a).

RSA has been used to discriminate between *animate* and *inanimate* entities with fMRI (Kriegeskorte et al., 2008b) and with MEG/EEG (Cichy et al., 2014; Cichy and Pantazis, 2017). MEG/EEG activity, measured at the scalp surface, contains rich spatial information about underlying representationally-specific patterns of neural activity, and it has the temporal resolution to track how similarities amongst these patterns change over time (Stokes et al., 2015). Here, we used RSA in combination with MEG and EEG to ask whether comprehenders can use the animacy constraints of verbs to predict the semantic features associated with the animacy of upcoming nouns. If this is the case, the similarity in spatial patterns should be greater following *animate constraining* than *inanimate constraining* verbs, reflecting the greater intercorrelation amongst predicted animate than predicted inanimate semantic features of the upcoming noun. Moreover, if these animacy predictions are generated irrespective of being able to predict specific words, this effect should be equally large following *low discourse constraint* and *high discourse constraint* contexts.

## Materials & Methods

### Overall structure of experiments and analysis approach

We carried out two studies using the same experimental design and overlapping sets of stimuli. In the first study, we collected MEG and EEG data simultaneously in 32 participants. In the second study, we collected EEG data in 40 different participants. We analyzed the MEG data and the EEG data separately. For the EEG analysis, we used the EEG data from participants in both the first and second studies to maximize statistical power (n=72).

In this Methods section, we first introduce the experimental design and stimuli, which were used in both the first MEG-EEG study and the second EEG-only study. Second, we describe the participants and overall procedures in each of the two studies. Third, we report MEG data acquisition and preprocessing (for the first MEG-EEG study), and EEG data acquisition and preprocessing (for both the first MEG-EEG study and the second EEG-only study). Fourth, we describe the spatial similarity analysis, which was the same for the MEG and the EEG datasets. We also describe an analysis of the evoked responses produced by the verb — event-related fields (ERFs) for the MEG data and ERPs for the EEG data — which was carried out to constrain our interpretation of the spatial similarity findings.

### Experimental design and stimuli

#### Experimental design

In both the MEG-EEG study and the EEG-only study, stimuli were three-sentence scenarios (Table 1). The first two sentences introduced a discourse context, and the final sentence began with an adjunct phrase of 1-4 words, followed by a pronominal subject that referred back to a protagonist introduced in the first two sentences, followed by a verb. Following the verb, there was a determiner, a direct object noun, and then three additional words to complete the sentence. The verb in the third sentence, which we refer to as the *critical verb*, varied in whether it constrained for an *animate* direct object noun (*animate constraining:* 50%, e.g. “cautioned the…”) or an *inanimate* direct object noun (*inanimate constraining:* 50%, e.g. “unfolded the…”). In addition, the lexical constraint of full discourse context (the combination of the first two sentences and the first few words of the third sentence, including the verb and the determiner) varied such that it predicted a single word (*high discourse constraint*: 50%, e.g. “The lifeguards received a report of sharks right near the beach. Their immediate concern was to prevent any incidents in the sea. Hence, they cautioned the…”), or for no specific single word (*low discourse constraint*, e.g. “Eric and Grant received the news late in the day. They mulled over the information, and decided it was better to act sooner rather than later. Hence, they cautioned the…”). This crossing of Verb animacy constraint (*animate constraining, inanimate constraining*), and Discourse constraint (*high discourse constraint*, *low discourse constraint*) gave rise to the four conditions relevant to the present study.

**Table 1.** Examples of the four experimental conditions.

| Verb animacy constraint | Discourse constraint | Example | *Lexical constraint of discourse context |
| --- | --- | --- | --- |
| Animate constraining | High discourse constraint | The lifeguards received a report of sharks right near the beach. Their immediate concern was to prevent any incidents in the sea. Hence, they <u>cautioned</u> the ... | 65% (15%) |
|  | Low discourse constraint | Eric and Grant received the news late in the day. They mulled over the information, and decided it was better to act sooner rather than later. Hence, they <u>cautioned</u> the ... | 19% (11%) |
| Inanimate constraining | High discourse constraint | Judith was working on the origami project for her office fundraiser. She was starting to get frustrated because it was her third attempt at making a crane. Nevertheless, she <u>unfolded</u> the ... | 71% (14%) |
|  | Low discourse constraint | Judith was nearing the end of her rope. She didn't think she could keep going. Nevertheless, she <u>unfolded</u> the ... | 24% (13%) |

Note: Discourse scenarios were created around *animate constraining* and *inanimate constraining* verbs (“cautioned” and “unfolded”; underlined here but not in the experiment itself). The sentences continued with object nouns plus three additional words, as indicated by the three dots.
\*The lexical constraint of the discourse context was operationalized as the percentage of participants who produced the best completion in a cloze study (see main text). Mean values are shown with standard deviations in parentheses.

Following the verb, the direct object noun either confirmed (e.g. “trainees”) or violated (e.g. “drawers”) the verb’s animacy constraint, rendering the scenarios either plausible or anomalous. In this manuscript, however, our focus was on the neural activity associated with the prediction of the upcoming noun, and so we report activity following the onset of the verb until just *before* the onset of the direct object noun. A full analysis of activity produced by the following nouns in both the MEG and EEG datasets (spatial similarity patterns as well as evoked responses), together with a detailed discussion of the relationships between these measures, will be reported in a separate paper (Wang and Kuperberg, Unpublished).

#### Construction of scenarios

In order to construct these scenarios, we began with a large set of preferentially transitive verbs. We established their animacy constraints as well as their lexical constraints in minimal contexts by carrying out an offline cloze norming study, described below. Then, on the basis of these norms, we selected a subset of *animate* and *inanimate* constraining verbs, which, in these minimal contexts, did not constrain strongly for a specific upcoming noun. For each verb, we then wrote discourse scenarios, and for each scenario, we quantified the constraints of the entire discourse context (the first two sentences plus the third sentence until after the determiner) with a second cloze norming study, described below.

##### Cloze norming studies

In both cloze norming studies, participants were recruited through Amazon Mechanical Turk. They were asked to complete each context with the first word that came to mind (Taylor, 1953) and, in an extension of the standard cloze procedure, to then provide two additional words that could complete the sentence (see Schwanenflugel and LaCount, 1988; Federmeier et al., 2007). Responses were excluded in any participant who indicated that the first language they learned was anything other than English, or if they reported any psychiatric or neurological disorders. Responses were also excluded in any participants who failed to follow instructions (“catch” questions were used as periodic attention checks).

###### Cloze norming study 1: To select a set of verbs based on their animacy and lexical constraints in minimal contexts

We began with a set of 617 transitively-biased verbs, compiled from various sources including Levin (1993) and materials from previous studies conducted in our laboratory (Paczynski and Kuperberg, 2011, 2012). Verbs with log Hyperspace Analogue to Language (HAL) frequency (Lund and Burgess, 1996) of two standard deviations below the mean (based on English Lexicon Project database: Balota et al., 2007) were excluded. For each verb, we constructed a simple active, past tense sentence stem that consisted of only a proper name, the verb, and a determiner (e.g., “Harry cautioned the…”). These sentences were divided into six lists in order to decrease the time demands on any individual participant during cloze norming. Between 89 and 106 participants (depending on list) who met inclusionary criteria provided completions for each verb.

For each verb, we identified the best completion of the sentence context (i.e. the most common first noun produced across all participants), and, based on the animacy of these nouns, we categorized the verb as either *animate constraining* or *inanimate constraining*. We also tallied the number of participants who produced this best completion in order to calculate the lexical constraint of the verbs for specific upcoming nouns in these minimal contexts. To generate the final set of discourse stimuli, we selected 175 verbs (88 *animate constraining* and 87 *inanimate constraining*), all with lexical constraints of lower than 24%.

###### Cloze norming study 2: To establish the constraint of the entire discourse contexts for upcoming nouns

For each of the *animate constraining* and *inanimate constraining* verbs, we wrote two types of two-sentence contexts. These contexts, in combination with the first few words of the third sentence, the verb, and the determiner, aimed either to constrain for either a single upcoming word (*high discourse constraint)* or for multiple possible upcoming words (*low discourse constraint)*. We then carried out a second cloze norming study of these discourse contexts to quantify their discourse constraints. The *high discourse constraint* and *low discourse constraint* contexts were pseudorandomly divided into two lists such that each list contained only one of the two types of discourse contexts associated with each verb. The two lists were then divided into thirds to decrease time demands on any individual participant during cloze norming. Between 51 and 69 participants who met inclusionary criteria provided completions for each scenario.

We found that, following both the *animate constraining* and the *inanimate constraining* verbs, over 99% of the completions produced were nouns. Similar to the first cloze norming study, the lexical constraint of each discourse context was calculated by tallying the number of participants who produced the most common completion in each discourse context. The mean lexical constraint of the *high discourse constraint* contexts was 67.80% (SD: 15.00%) and the mean lexical constraint of the *low discourse constraint* context was 21.56% (SD: 12.00%), and this differed significantly between the two conditions, t_(698)_ = 45.01, *p* < .001.

#### Distribution of stimuli into lists

The stimuli were then divided into lists, with each list containing (approximately) 50% *animate constraining* verbs and 50% *inanimate constraining* verbs, distributed evenly across the *high discourse constraint* and the *low discourse constraint* contexts. The lists were constructed so that the same verb was not combined with the same discourse context more than once, but so that, across lists, all critical verbs were combined with both *high discourse constraint* and *low discourse constraint* contexts. Although the present study focuses on activity prior to the onset of the direct object noun, we constructed scenarios so that the subsequent direct object noun either confirmed the animacy constraints of the verb (and so the scenario was plausible) or violated the animacy constraints of the verb (and so the scenario was anomalous). The lists were constructed so that each participant viewed 50% plausible scenarios (one quarter of these plausible scenarios contained lexically predictable nouns following *high discourse constraint* contexts), and 50% anomalous scenarios. Thus, a scenario was just as likely to be plausible following a *high discourse constraint* context as following a *low discourse constraint* context.

In the first MEG-EEG study, the stimuli constituted 700 scenarios, which were divided into four lists, with each list containing 200 scenarios. Within each list, 101 scenarios contained *animate constraining* verbs and 99 scenarios contained *inanimate constraining* verbs. Since there were 175 unique verbs in total (88 *animate constraining* and 87 *inanimate constraining*), this meant that a small number of verbs in the third sentence were repeated: 13 out of 101 *animate constraining* verbs and 12 out of 99 *inanimate constraining* verbs.

In the second EEG-only study, we included a subset of 600 scenarios, which were divided into five lists. Each list contained 160 scenarios, with no verb being repeated in any of the lists (80 unique *animate constraining* and 80 unique *inanimate constraining* verbs). A detailed description of the precise counterbalancing scheme can be found in Kuperberg et al. (2019).

#### Quantification of the semantic and lexical similarity structures of the verbs

##### Semantic similarity structure of the animate constraining and the inanimate constraining verbs

In order to be able to infer that any difference in the similarity of the spatial pattern of neural activity following the *animate* and *inanimate constraining* verbs was due to the prediction of animacy features associated with the upcoming nouns for which they constrained, it was important to verify that the two groups of verbs did not differ markedly in other aspects of their internal semantic similarity structure. In particular, it was important to verify that the *animate constraining* verbs were not more similar to each other than the *inanimate constraining* verbs. Of course, some aspects of verb meaning are inherently tied to the meaning of the arguments for which they constrain (McCawley, 1968; Jackendoff, 1993), and the goal of the present study was to ask whether these types of constraints were used to *predict* upcoming animacy features as the sentences unfolded in real time. However, many other aspects of a verb’s meaning are not directly linked to the meaning of their arguments, and it was important to check that these other features of the verbs didn’t covary with their animacy constraints. For example, the two *animate constraining* verbs, “cautioned” and “alarmed”, are more similar to each other than the two *inanimate constraining* verbs, “folded” and “distributed”, not *only* because they both constrain for upcoming animate features, but *also* because both their meanings are specific instances of the broad meaning of “warn”.

In order to quantify these other components of verb meaning, we used WordNet, an English lexical database that groups words together based on their semantic relations (Miller et al., 1990), and that has been integrated in the Natural Language Toolkit (NLTK) (Loper and Bird, 2002). In WordNet, verbs are organized into hierarchies based on their semantic relations (Fellbaum, 1990), such as specificity in manner (e.g. walking – strolling), entailments (e.g. snoring – sleeping), causation (e.g. drop – break) and antonymy (e.g. coming – going). By examining the hierarchical structure of this network, the semantic similarity between different verbs can be quantified.

When examining the WordNet hierarchy for a given word, it is important to first consider its precise meaning in context — its so-called *sense*. For instance, the verb, “caution”, has at least two senses, including (a) “warn strongly; put on guard”, denoted in WordNet as Synset(‘caution.v.01’), and (b) “the trait of being cautious; being attentive to possible danger”, denoted as Synset(‘caution.n.01’). Therefore, for each of our critical verbs, a native English speaker manually identified its sense within each discourse context. For example, the sense of the verb “cautioned” within the example scenario shown in Table 1 (“The lifeguards received a report of sharks right near the beach. Their immediate concern was to prevent any incidents in the sea. Hence, they cautioned …”) was classified as Synset(‘caution.v.01’). In total, across the entire stimulus set, we identified 250 unique verb senses (113 *animate constraining*, 137 *inanimate constraining*).

We then calculated semantic similarity values between all possible pairs of verb senses within the sets of *animate constraining* and *inanimate constraining* verbs. As a measure of semantic similarity, we used a path-based approach described by Wu & Palmer (Wu and Palmer, 1994), which is known to correlate with human ratings of semantic similarity (Slimani, 2013). Wu & Palmer similarity values range between 0 and 1, with values approaching 0 indicating low similarity, and a value of 1 indicating identical concepts. We stored these pairwise Wu-Palmer semantic similarity values in a 250 by 250 symmetric semantic similarity matrix, with rows and columns indexing the individual verbs’ senses, see Figure 1A. Examination of this matrix did not reveal any clear difference in the internal semantic similarity structure between the *animate constraining* verbs (top-left: semantic similarity values for verb senses, 1 to 113: 113*112/2 = 6328 pairs) and the *inanimate constraining* verbs (bottom-right: semantic similarity values for verb senses, 114 to 250: 137*136/2 = 9316 pairs).

**Figure 1.**
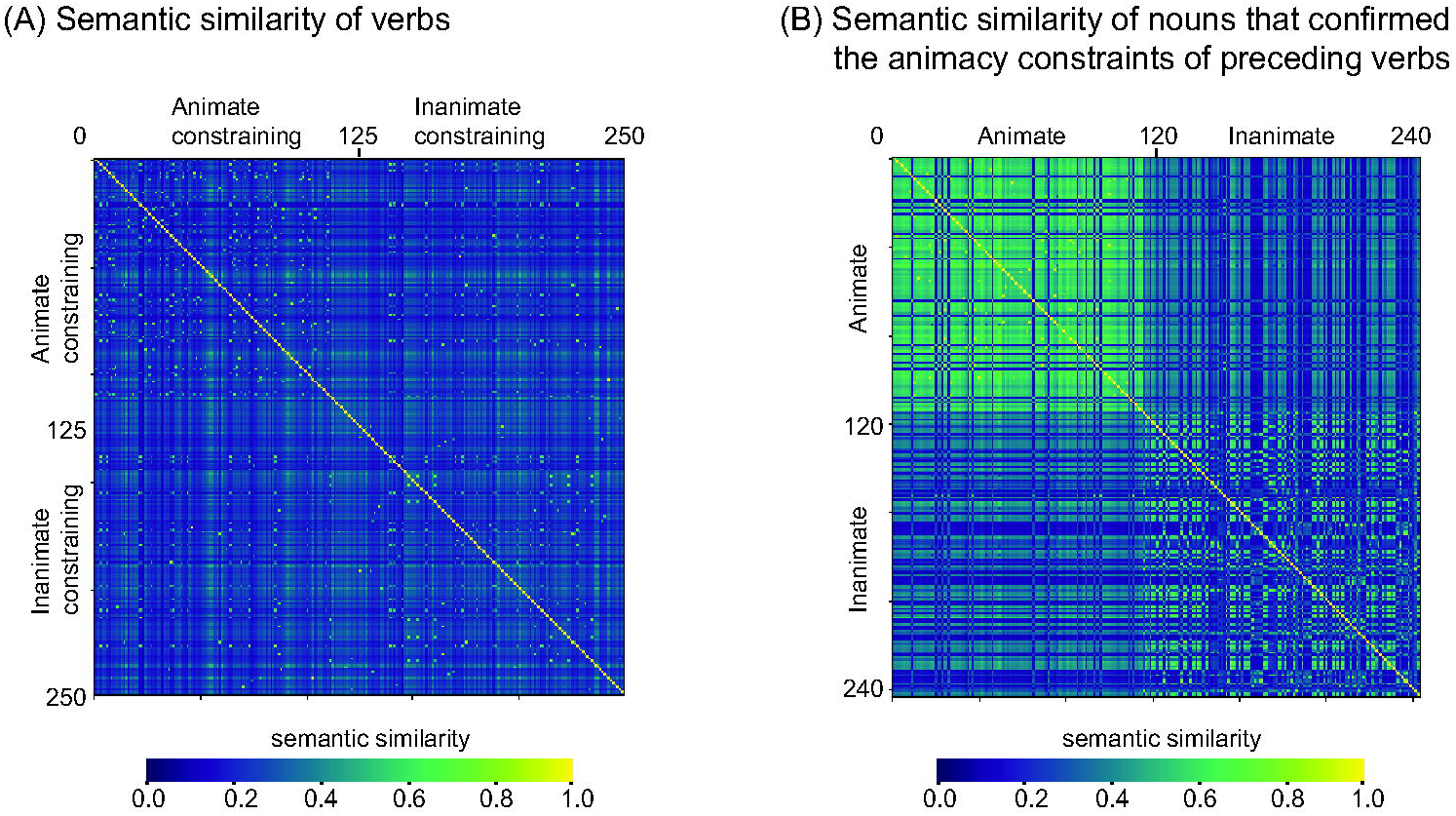
Pairwise Wu & Palmer similarity values for the senses of (A) *animate constraining* and *inanimate constraining* verbs, and (B) nouns that confirmed these animacy constraints. The range of these Wu & Palmer similarity values is between 0 and 1, with values approaching 0 indicating low similarity, and a value of 1 indicating identical concepts. (A) Pairwise Wu-Palmer semantic similarity values of the verbs are shown in a 250 by 250 symmetric semantic similarity matrix, with rows and columns indexing the individual verbs’ senses (*animate constraining* verbs: from 1 to 113; *inanimate constraining* verbs: from 114 to 250). The pairwise Wu-Palmer semantic similarity values of the *animate constraining* verbs (values at the top-left of the matrix) were smaller than those of the *inanimate constraining* verbs (values at the bottom-right of the matrix): *p* = .04 (1000 permutations). (B) Pairwise Wu-Palmer semantic similarity values of the nouns are shown in a 244 by 244 symmetric semantic similarity matrix, with rows and columns indexing the individual nouns’ senses (animate nouns: from 1 to 116; inanimate nouns: from 117 to 244). The pairwise Wu-Palmer semantic similarity values of the *animate nouns* (values at the top-left of the matrix) were larger than those of the *inanimate nouns* (values at the bottom-right of the matrix): *p* = .001 (1000 permutations).

To test this statistically, we carried out a permutation-based statistical test on these pairwise similarity values, after excluding the values of 1s along the diagonal line. We extracted the Wu & Palmer semantic similarity values for each possible pair of *animate constraining* verbs (113*112/2 = 6328 values) and took the mean value, and we did the same for each possible pair of *inanimate constraining* verbs (137*136/2 = 9316 values). We then took the difference in these means as our test statistic. After that, we randomly re-assigned the similarity values across the two groups of verbs, and re-calculated the mean difference between the two groups. We took the mean difference value for each randomization (1000 times) to build a null distribution. If the observed test statistic fell within the highest or lowest 2.5% of this distribution, it was considered to be significant. This test showed that the semantic similarity among the *animate constraining* verbs (mean +/− SD = 0.24 +/− 0.09) was very slightly *lower* than that among the *inanimate constraining* verbs (mean +/− SD = 0.26 +/− 0.08), *p* = .04.

##### Lexical similarity structure of the animate constraining and the inanimate constraining verbs

We also verified that the two groups of verbs did not differ in various aspects of their internal *lexical* similarity structures. To do this, we extracted the following lexical properties of each verb: length (i.e. number of letters), orthographic Levenshtein distance (OLD20, Balota et al., 2007) and log frequency (based on the SUBTLEX database Brysbaert and New, 2009). For each of these lexical variables, we calculated the absolute difference for each possible pair of *animate constraining* (88*87/2 = 3828 values) and *inanimate constraining* verbs (87*86/2 = 3741 values). As described above, we then calculated the mean value in each group and took the difference in these means as our test statistic, and tested for differences in the lexical similarity structure between the two groups of verbs using a permutation test (1000 permutations). This test showed that the internal similarity structures, based on length, orthographic neighborhood and the frequency, were matched between the *animate constraining* and *inanimate constraining* verbs, all *p*s > .07.

#### Quantification of the semantic and lexical similarity structures of the predicted nouns

##### Semantic similarity structure of animate and inanimate nouns constrained for by the verbs

Our main hypothesis rested on the assumption that the predicted *animate* nouns would be more semantically similar to each other than the predicted *inanimate* nouns. Obviously, we had no way of knowing precisely what nouns each participant would predict during the experiment itself, particularly in the low constraint discourse contexts. Therefore, as proxy for these semantic predictions, we took 350 (50%) of the *animate* and *inanimate* direct object nouns that participants actually viewed — those that confirmed the animacy constraints of the verbs, rendering the scenarios plausible — and quantified their semantic similarity structures. We again used WordNet in which the meaning relationships of nouns, such as super-subordinate relations (e.g. furniture – chair) and part-whole relations (e.g. chair – backrest), are organized in a hierarchical network (Miller, 1990). We again quantified their semantic similarity using Wu-Palmer semantic similarity values (Wu and Palmer, 1994).

Just as described above for the verbs, we first manually identified the sense of each noun within its preceding context for all 350 plausible scenarios (175 *animate nouns*, 175 *inanimate nouns*), resulting in 244 unique senses for the nouns (116 *animate nouns*, 128 *inanimate nouns*). We stored the Wu-Palmer semantic similarity values (Wu and Palmer, 1994) for all possible pairs of the nouns’ senses in a 244 by 244 matrix, see Figure 1B. The similarity values shown at the top-left of the matrix represent the pairwise Wu-Palmer semantic similarity values for all pairs of *animate nouns* (nouns 1 to 116: 116*115/2 = 6670 pairs), and the similarity values shown at the bottom-right of the matrix represent the pairwise semantic similarity values for all pairs of *inanimate nouns* (nouns 117 to 244: 128*127/2 = 8128 pairs). This matrix suggests that the pairwise Wu-Palmer semantic similarity values for the *animate nouns* were indeed larger than those for the *inanimate nouns*. A permutation-based statistical test (1000 permutations, carried out as described above) confirmed this observation (*animate nouns*: mean +/− SD = 0.49 +/− 0.20; *inanimate nouns:* mean +/− SD = 0.29 +/− 0.19, *p* = .001).

##### Lexical similarity structure of the animate and inanimate nouns constrained for by their preceding verbs

Finally, it was important to check that any differences in similarity between the neural patterns of activity produced by predicted *animate* and *inanimate* nouns were not driven by differences in the similarity of their lexical features rather than of their semantic features. Again, we had no way of knowing precisely what nouns each participant would predict during the experiment itself. However, we knew that 100 scenarios had *high discourse constraints* and were followed by predicted nouns. Therefore, as a proxy for any lexical-level predictions, we extracted the lexical properties of the predicted nouns that followed these *high discourse constraint* contexts (the most commonly produced nouns in the second cloze norming study described above): length (number of letters), orthographic Levenshtein distance (OLD20, Balota et al., 2007) and log frequency (Brysbaert and New, 2009). For each of these variables, we again calculated the absolute difference values between each possible pair of predicted *animate* nouns (50*49/2 = 1225 values) and predicted *inanimate* nouns (50*49/2 = 1225 values). Then we calculated the mean value in each group and took the difference as our test statistic, and tested for any difference in the lexical similarity structure between the two groups of nouns using the permutation test described above (1000 permutations). This test revealed no statistically significant differences for word length, frequency or orthographic neighborhood (all *p*s > .15).

### Participants

The first MEG-EEG dataset was acquired at Massachusetts General Hospital. Written consent was obtained from all participants following the guidelines of the Massachusetts General Hospital Institutional Review Board. Thirty-three participants initially participated, but we subsequently excluded the data of one participant because of technical problems. This left a final dataset of 32 participants (16 females, mean age: 23.4 years; range 18-35 years).

The second EEG-only dataset was acquired at Tufts university. Participants gave informed consent following procedures approved by the Tufts University Social, Behavioral, and Educational Research Institutional Review Board. Data were collected from 40 participants (19 females, mean age: 21.5 years; range 18-32 years).

In both experiments, all participants were right-handed as assessed using the modified Edinburgh Handedness Inventory (Oldfield, 1971; White and Ashton, 1976). All had normal or corrected-to-normal vision and were native speakers of English with no additional language exposure before the age of 5. Participants were not taking psychoactive medication, and were screened to exclude the presence of psychiatric and neurological disorders.

### Overall procedure

In both studies, stimuli were presented using PsychoPy 1.83 software (Peirce, 2007) and projected on to a screen in white Arial font on a black background, with a size that was one-tenth of the screen height. The first two sentences were each presented as a whole (each for 3900ms, 100ms interstimulus interval, ISI), followed by an intra-trial fixation (white “++++”), which was presented for 550ms, followed by a 100ms ISI. The third sentence, which contained the *animate constraining* or *inanimate constraining* verb, was presented word by word (each word for 450ms, 100ms ISI). The final word of the third sentence was followed by a pink “?” (1400ms, 100ms ISI). This cued participants to press one of two buttons with their left hand to indicate whether each discourse scenario “made sense” or not (response fingers were counterbalanced across participants). In addition, after a proportion of trials (24/200 in the MEG-EEG study; 32/160 in the EEG-only study; semi-randomly distributed across runs), a comprehension question, referring to the immediately previous scenario, appeared on the screen (1900ms, 100ms ISI). Participants were asked to respond yes or no based on the scenario they just read. This encouraged them to attend to and comprehend the scenarios as a whole, rather than focusing only on the third sentence. Following each trial, a blank screen was presented with a variable duration that ranged from 100 to 500ms. This was then followed by a green fixation (++++) for a duration of 900ms followed by an ISI of 100ms. Participants were encouraged to blink during the green fixation period.

In both studies, stimuli were presented over several runs (in the MEG-EEG study, 200 scenarios presented over eight runs, each with 25 scenarios; in the EEG-only study, 160 scenarios presented over four runs, each with 40 scenarios). Runs were presented in random order in each participant. Before the onset of each study, a practice session was conducted to familiarize participants with the stimulus presentation and the judgment tasks.

#### MEG data acquisition and preprocessing

##### MEG data acquisition

In the first MEG-EEG study, MEG data were acquired together with EEG data (the EEG setup is described below). Participants sat inside a magnetically shielded room (IMEDCO AG, Switzerland), and MEG data were acquired with a Neuromag VectorView system (Elekta-Neuromag Oy, Finland) with 306 sensors (102 triplets, each comprising two orthogonal planar gradiometers and one magnetometer). Signals were digitized at 1000Hz, with an online bandpass filter of 0.03 - 300Hz. To monitor for blinks and eye movements, Electrooculography (EOG) data were collected with bipolar recordings: vertical EOG electrodes were placed above and below the left eye, and horizontal EOG electrodes were placed on the outer canthus of each eye. To monitor for cardiac artifact, electrocardiogram (ECG) data were collected, also with bipolar recordings: ECG electrodes were placed a few centimeters under the left and right collarbones. At both EOG and ECG sites, impedances were kept at less than 30 kΩ. To record the head position relative to the MEG sensor array, the locations of three fiduciary points (nasion and two auricular), four head position indicator coils, all EEG electrodes, and at least 100 additional points were digitized using a 3Space Fastrak Polhemus digitizer, integrated with the Vectorview system. Before each run, we used the four head position indicator coils to monitor the position and orientation of the head with respect to the MEG sensor array.

##### MEG data preprocessing

MEG data were preprocessed using version 2.7.4 of the Minimum Norms Estimates (MNE) software package in Python (Gramfort et al., 2014). In each participant, in each run, MEG sensors with excessive noise were visually identified and removed from further analysis. This resulted in the removal of seven (on average) out of the 306 MEG sensors. Eye-movement and blink artifacts were automatically removed using the algorithms recommended by Gramfort et al. (2013). Signal-Space Projection (SSP) correction (Uusitalo and Ilmoniemi, 1997) was used to correct for ECG artifact. Then, after applying a bandpass filter at 0.1 to 30Hz, we segmented data into -100 to 2100ms epochs (relative to verb onset). Epochs in which the range of amplitudes exceeded pre-specified cutoff thresholds (4e-10 T/m for gradiometers and 4e-12 T for magnetometers) were removed. The data of bad MEG sensors were interpolated using spherical spline interpolation (Perrin et al., 1989). Our epoch of interest for analysis was from -100 to 1100ms, relative to verb onset. On average, 85 artifact-free trials remained following the *animate constraining* verbs and 83 trials remained following the *inanimate constraining* verbs, with no statistically significant difference between the two groups: F_(1,31)_ = 3.94, *p* = .06, η^2^ = 0.11.

#### EEG data acquisition and preprocessing

##### EEG data acquisition

The first EEG dataset was acquired simultaneously with the MEG data using a 70-electrode MEG-compatible scalp electrode system (BrainProducts, München). The EEG signals were digitized at 1000Hz, with an online bandpass filter of 0.03 - 300Hz. The second EEG dataset was recorded using a Biosemi Active-Two acquisition system from 32 active electrodes in a modified 10/20 system montage. Signals were digitized at 512Hz, with a bandpass of DC - 104Hz, and EEG electrodes were referenced offline to the average of the left and right mastoid electrodes. Impedances were kept at < 30kΩ at all scalp sites for both studies.

##### EEG data preprocessing

Both EEG datasets were preprocessed using the Fieldtrip software package, an open-source Matlab toolbox (Oostenveld et al., 2011). For spatial similarity analysis, we planned to combine all participants in the two EEG datasets to maximize power. Therefore, given that the two datasets were acquired with different online filtering settings (0.03 - 300Hz vs. DC - 104Hz), we applied an offline low-pass filter of 30Hz to the first EEG dataset, and an offline band-pass filter of 0.1 - 30Hz to the second EEG dataset. In addition, because the two datasets were acquired with different sampling rates (1000Hz vs. 512Hz), we down-sampled both datasets to 500Hz.

Each individual’s EEG data was segmented into epochs. We identified and removed, on average, seven bad EEG electrodes out of the 70 electrodes in the first EEG dataset, whereas no bad electrodes were identified or removed in the second EEG dataset. We applied an Independent Component Analysis (ICA; Bell and Sejnowski, 1997; Jung et al., 2000) and removed components associated with eye movement from the EEG signal. We then inspected the data visually and removed any remaining artifacts. The data of the seven bad EEG electrodes in the first dataset were then interpolated using spherical spline interpolation (Perrin et al., 1989).

On average, slightly more artifact-free trials remained following the *animate constraining* (81 trials on average) than the *inanimate constraining* verbs (79 trials on average), F_(1,71)_ = 9.12, *p* = .004, η^2^ = 0.114.

#### Spatial similarity analysis of both MEG and EEG data

We used the same method of carrying out the spatial similarity analysis for the MEG and the EEG data, using MATLAB 2014b (MathWorks) with custom-written scripts. A schematic illustration of our spatial similarity stream is shown in Figure 2. Note that this approach is somewhat different from classic RSA streams, which ask the question of whether dissimilarities amongst items along a particular dimension (in the present study, animacy) can be used to *discriminate* dissimilarity patterns of neural activity based on that dimension (Kriegeskorte et al., 2008a). For this study, we were not only interested in whether and when it was possible to discriminate between predicted animate and inanimate nouns based on neural activity; we were also interested in whether the similarity amongst patterns of brain activity associated with predicted animate nouns was greater than the similarity amongst patterns of activity associated with predicted inanimate nouns. We therefore computed average spatial similarity values at each time point following the *animate constraining* and the *inanimate constraining* verbs separately in each individual participant. Specifically, in each participant, for each trial, and at each time point, we extracted a vector of data that represented the spatial pattern of neural activity across all channels (MEG: 306 sensors; EEG dataset from the first study: 70 electrodes; EEG dataset from the second study: 32 electrodes). At each time point, *t*, we quantified the similarity between the spatial pattern of neural activity following all possible pairs of *animate constraining* verbs (e.g. between A-S1 and A-S2, between A-S1 and A-Sn, in Figure 2) and all possible pairs of *inanimate constraining* verbs (e.g. between B-S1 and B-S2, between B-S1 and B-Sn in Figure 2) by calculating a *Pearson’s r* value between the spatial vectors. These pairwise correlation R-values were used to construct spatial similarity matrices at each time point, corresponding to the spatial similarity patterns of neural activity following the *animate constraining* verbs (the left matrix in Figure 2) and the *inanimate constraining* verbs (the right matrix in Figure 2). We then averaged the N*(N-1)/2 off-diagonal elements of these matrices to compute an averaged R-value at each time point in each participant.

**Figure 2.**
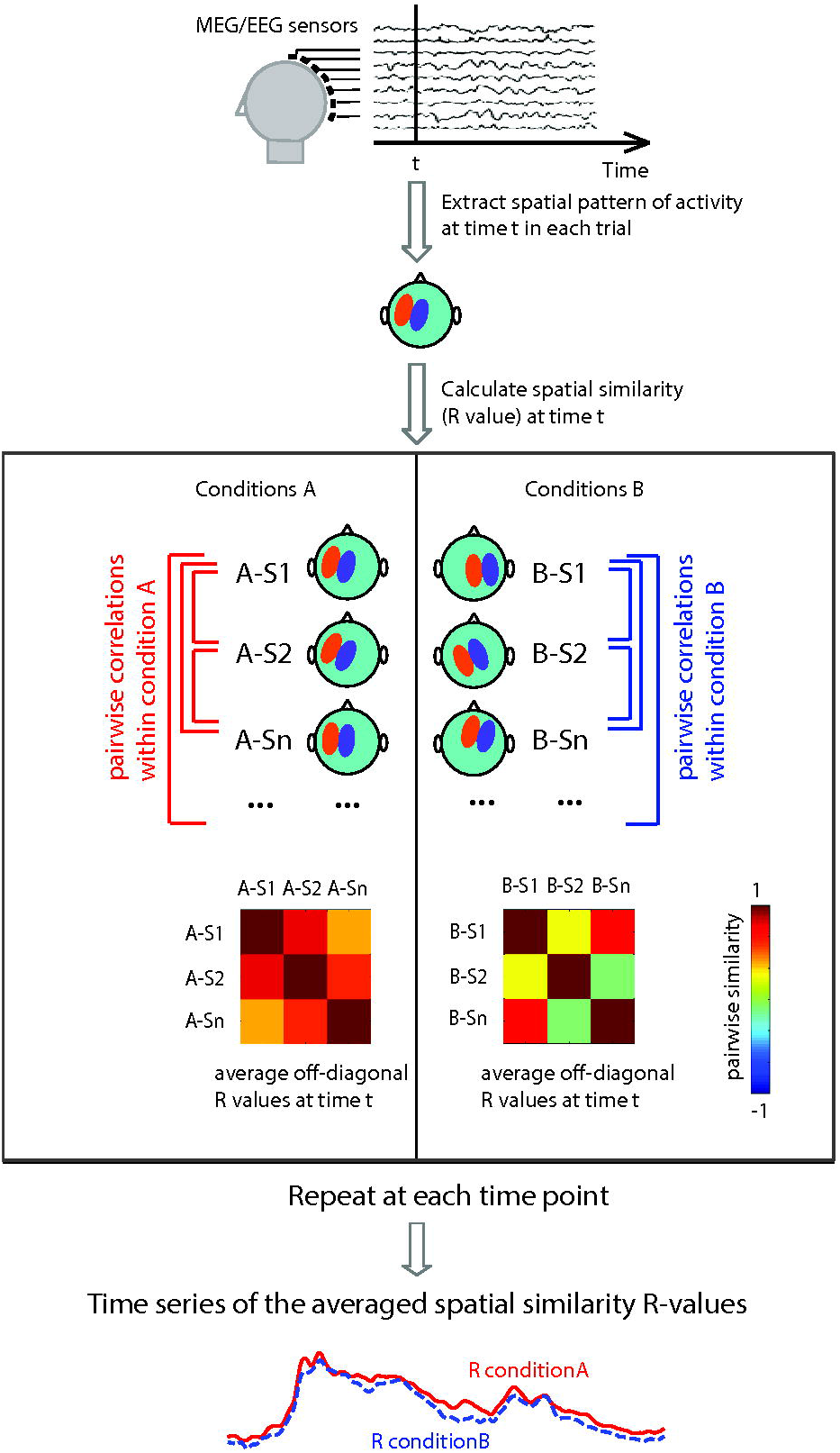
A schematic illustration of the spatial Representational Similarity Analysis in the present study. First, in each participant, for each trial, and at each time point, *t*, a vector of data was extracted across all MEG/EEG sites to represent the spatial pattern of neural activity produced at that time point. Second, at each time point, *t*, the similarity between the spatial pattern of neural activity produced by all possible pairs of trials of condition A (e.g. between A-S1 and A-S2, between A-S1 and A-Sn within the left side of the box) and all possible pairs of trials of condition B (e.g. between B-S1 and B-S2, between B-S1 and B-Sn within the right side of the box) was quantified by calculating the *Pearson’s r* values between the spatial vectors. These pairwise correlation R-values were used to construct spatial similarity matrices at each time point, corresponding to the spatial similarity patterns of neural activity produced in condition A (the left matrix) and condition B (the right matrix). Third, the N*(N-1)/2 off-diagonal elements of these matrices were averaged to compute an averaged R-value at each time point that corresponded to the average spatial similarity pattern produced by each of the two conditions. These average values at each consecutive time point yielded two time-series of spatial similarity R-values in each participant, reflecting the temporal dynamics of the spatial similarity of brain activity produced in conditions A (red solid line) and condition B (blue dotted line).

To visualize our data, we averaged these similarity values across all participants in the MEG dataset (n=32) and across all participants in the two EEG datasets (n=72) at each consecutive time point following the *animate constraining* verbs (the solid red line in Figure 2) and the *inanimate constraining* verbs (the dotted blue line in Figure 2). This yielded a ‘grand-average’ spatial similarity time series for each condition, which allowed us to visualize the timing and directionality of any differences between the two conditions. This visualization is analogous to traditional visualizations of grand-average evoked responses, and so it also helped us to directly compare the time course of the spatial similarity values with the evoked responses produced by the verbs.

For statistical analysis, we took the average spatial similarity values in each participant as the dependent measure and asked whether and when there were significant differences in the spatial similarity patterns of neural activity following the *animate constraining* and the *inanimate constraining* verbs. For the EEG spatial similarity analysis, we used the spatial similarity values of all 72 individuals in both EEG datasets to increase statistical power (we subsequently tested whether the two EEG datasets showed a statistically significant difference for the reported effect, see Results). We used cluster-based permutation tests to control for multiple comparisons across multiple time points (Maris and Oostenveld, 2007). Specifically, at each time point from the onset of the verb (t = 0) until before the direct object noun actually appeared (t = 1100ms), we carried out a paired t-test (550 tests in total). Adjacent data points that exceeded a preset uncorrected *p*-value threshold of *.05* were considered temporal clusters. The individual t-statistics within each cluster were summed to yield a cluster-level test statistic — the cluster mass statistic. We then randomly re-assigned the spatial similarity R-values across the two conditions (i.e. *animate constraining* and *inanimate constraining* verbs) at each time point within each participant, and calculated cluster-level statistics as described above. This was repeated 10000 times. For each randomization, we took the largest cluster mass statistic (i.e. the summed t values), and, in this way, built a null distribution for the cluster mass statistic. We then compared our observed cluster-level test statistic against this null distribution. Any temporal clusters falling within the highest or lowest 2.5% of the distribution were considered significant.

#### Analysis of the evoked responses of both MEG and EEG data

In order to constrain our interpretation of the similarity values, and in particular to verify our assumption that any differences in spatial similarity following the *animate constraining* versus the *inanimate constraining* verbs reflected the pre-activation of predicted animacy features of the upcoming noun, rather than lexico-semantic processing of the verbs themselves, we examined the evoked responses produced by the two types of verbs. We carried out a classic ERF analysis on the MEG data and an ERP analysis on the EEG data using the Fieldtrip software package (Oostenveld et al., 2011). For both the ERF and ERP analyses, we time-locked responses to verb onset, using a -100 - 0ms baseline, and we calculated evoked responses separately for the *animate constraining* and *inanimate constraining* verbs, collapsed across the high and low constraint discourse contexts, at each site in each participant. For the MEG data analysis, we used data from only the gradiometer sensors, combining these two sensors at each site by calculating the root mean square of the values.

To test for any differences in the ERFs/ERPs evoked by the *animate constraining* and *inanimate constraining* verbs, we again used cluster-based permutation tests using the Fieldtrip software package (Oostenveld, et al., 2011) to account for multiple comparisons over time points and channels (Maris & Oostenveld, 2007). The tests followed the same steps as described above, except that we carried out dependent-samples t-tests at each time point within the full 0-1100ms time window at each of the 102 MEG sensor sites and at each of the 32 EEG electrode sites (those that were used in both the MEG-EEG and EEG-only studies). All spatially and temporally adjacent data samples that exceeded a preset uncorrected significance threshold of 5% were taken as a spatiotemporal cluster, and individual t-statistics within each cluster were summed to yield cluster-level test statistics. These cluster-level test statistics were then compared against the null distribution that was built based on 1000 randomizations. Any spatiotemporal clusters falling within the highest or lowest 2.5th percentile were considered significant. To quantify the temporal extent of any significant clusters, we identified the first and last time points that revealed significant effects on at least three channels.

## Results

### Spatial similarity results

#### Spatial similarity results of the MEG data

Figure 3A, left, shows the group-averaged (32 participants) MEG time series of spatial similarity values following the *animate constraining* and the *inanimate constraining* verbs. This reveals a sharp increase in the overall degree of spatial similarity beginning at ∼50ms after verb onset, peaking twice between 100 and 200ms, and then decreasing with a third relatively broader peak between 300 - 400ms following verb onset. After that, the spatial similarity values decrease throughout the duration of the verb. A similar, rapid increase in the spatial similarity values can be seen following the onset of the determiner, which followed the verb at 550ms. This peaked at ∼150ms and ∼225ms following determiner onset before gradually decreasing again. These overall increases in spatial similarity values are likely to reflect the MEG equivalents of the N1/P2 and N400 evoked responses produced by the verb, and the N1/P2 produced by the determiner (which did not produce a large N400, as shown in Figure 4).

**Figure 3.**
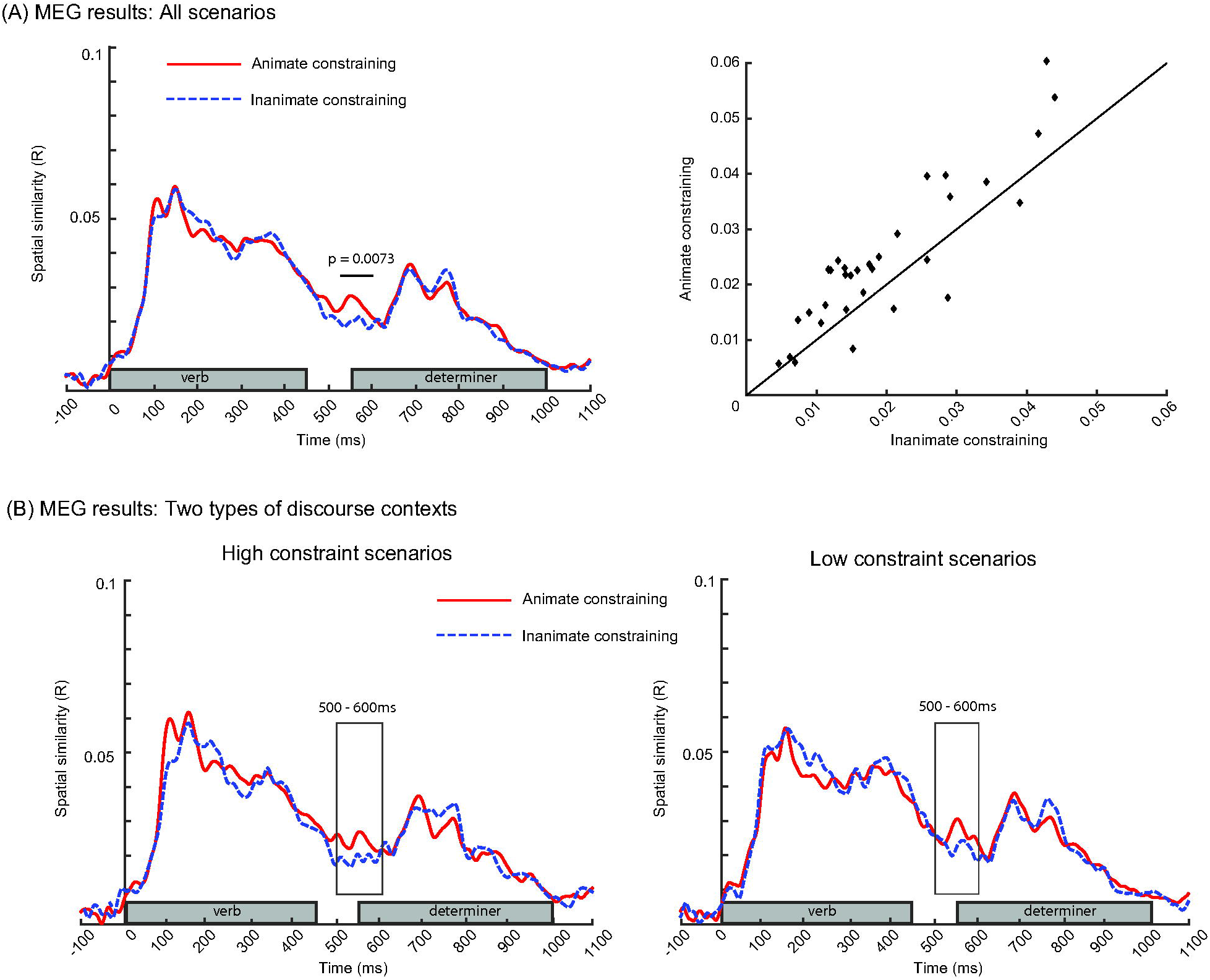
Results of the spatial similarity analysis of the MEG data (the first study, 32 participants). (A) Left: Group-averaged time series of spatial similarity values following *animate constraining* verbs (red solid line) and following *inanimate constraining* verbs (blue dotted line), from verb onset at 0ms to noun onset at 1100ms. The duration of the verbs (0 - 450ms) and the subsequent determiners (550 - 1000ms) are marked with grey bars on the x-axis. The spatial pattern of neural activity was more similar following the *animate constraining* than the *inanimate constraining* verbs between 529 - 599ms following verb onset (*p* = .0073, 10000 permutations); the significant cluster is highlighted by a black line over the time series. Right: A scatter plot of the averaged R-values per participant across the 500 - 600ms time window following verb onset. This shows that 26 of the 32 participants had R-values above the diagonal line, i.e. larger spatial similarity values following the *animate constraining* than the *inanimate constraining* verbs. (B) Group-averaged time series of the spatial similarity values following the *animate constraining* (red solid line) verbs and following the *inanimate constraining* verbs (blue dotted line) in the *high discourse constraint* scenarios that constrained strongly for a specific upcoming noun (left) and the *low discourse constraint* scenarios that did not constrain strongly for a specific noun (right). The spatial similarity effect was equally large following the two types of *discourse constraint* contexts, as indicated by the absence of an interaction between Verb animacy constraint and Discourse constraint (F(1,31) = 0.20, *p* = .66, η^2^ = 0.01), for the averaged spatial similarity values between 500 - 600ms following verb onset.

**Figure 4.**
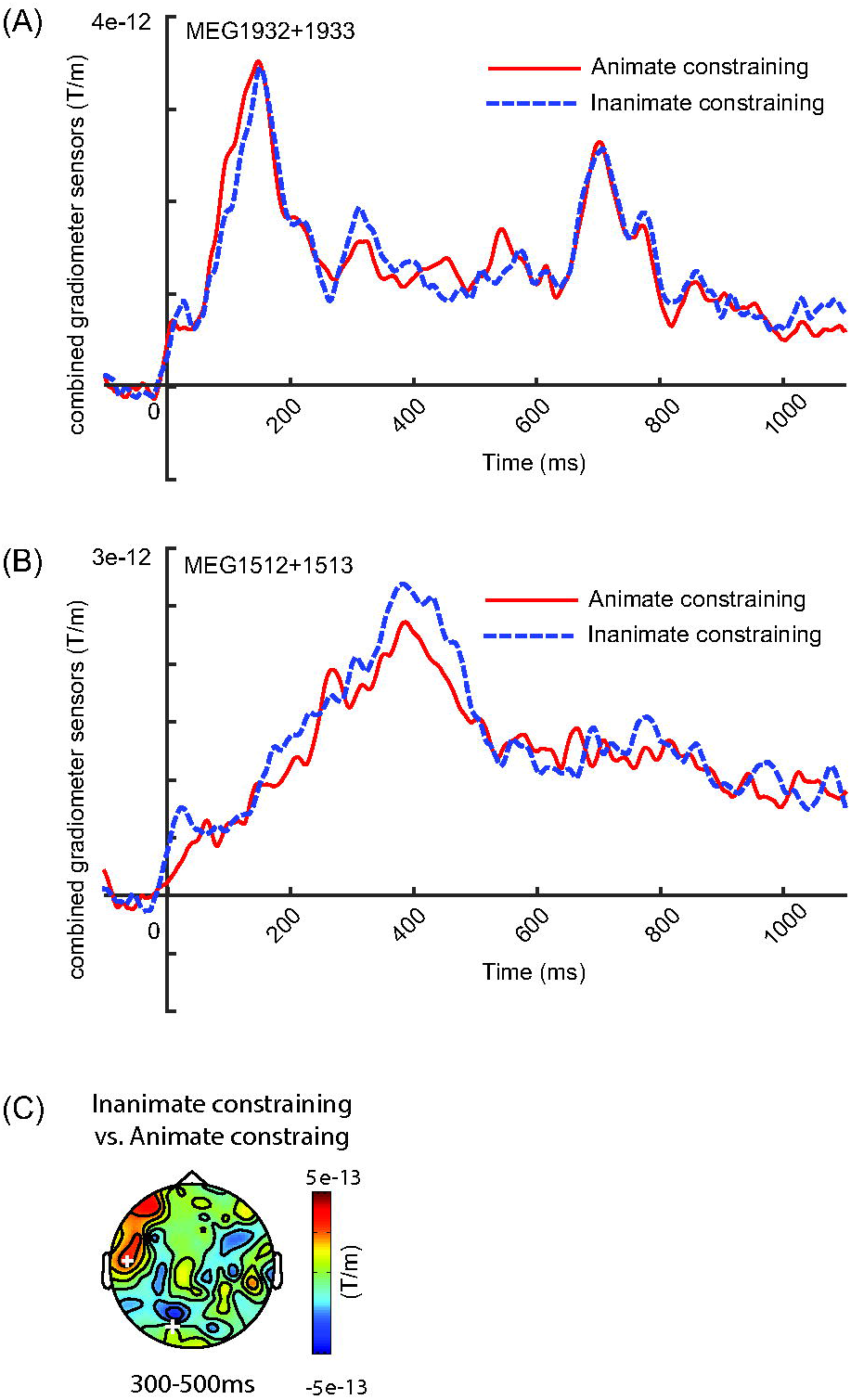
Grand-average event-related fields (ERFs) of the MEG data (n=32), time-locked to verb onset. ERFs following the onset of *animate constraining* and *inanimate constraining* verbs are shown at two representative MEG sensor sites (combined across two gradiometer sensors at each site) — (A) a left occipital sensor site (MEG1932+1933), and (B) a left temporal site (MEG1512+1513). Each of these sites is highlighted with a white cross on the topographic map (C). As shown in (A), following both the onset of the verb and the onset of the determiner (650 - 750ms after verb onset), the left occipital sensor shows clear stimulus-driven evoked responses between 100 - 200ms time window (the MEG equivalent of the N1/P2 component). As shown in (B), following the onset of the verb (but not the determiner), the left temporal sensor shows a strong evoked response between 300 - 500ms time window (the MEG equivalent of the N400 component). (C) The topographic distribution of the ERF difference within the 300 - 500ms time window. There was no significant ERF difference between the two conditions (*p* = .49) based on a cluster-based permutation test over the entire time window.

Of most relevance to the questions addressed in this study, from around the time of verb *offset* (450ms after verb onset), the spatial similarity patterns appeared to diverge such that the spatial patterns of neural activity were more similar following the *animate constraining* than the *inanimate constraining* verbs. This difference continued into the interstimulus interval (100ms), disappearing at ∼50ms following the onset of the determiner (i.e. lasting from ∼450 to ∼600ms after verb onset). A cluster-based permutation test (Maris and Oostenveld, 2007) across the entire epoch (0 - 1100ms) confirmed a significant difference in spatial similarity (*p* = .0073), with a cluster between 529 - 599ms following verb onset (although note that this is likely to underestimate of the true extent of the effect, see Maris and Oostenveld, 2007). As shown in Figure 3A, right, 26 of the 32 participants had larger spatial similarity values (averaged across the 500 - 600ms time window following verb onset) following the *animate constraining* than the *inanimate constraining* verbs.

We also compared the ERFs of the two conditions to determine whether the larger spatial similarities following the *animate constraining* versus *inanimate constraining* verbs could be explained by differences in the ERFs evoked by these verbs. As shown in Figure 4, if anything, the ERF evoked by the *inanimate constraining* verbs appeared to be larger than that evoked by the *animate constraining* verbs within the N400 time window, and a cluster-based permutation test over the entire epoch failed to reveal a significant ERF effect (*p* = .49).

We then asked whether this spatial similarity effect was modulated by overall discourse constraint — that is, whether it depended on being able to predict a specific upcoming lexical item. In minimal contexts, all verbs had relatively low lexical constraints (< 24%, as verified by our first cloze norming study). However, by design, and as verified by our second cloze norming study, 50% of the *animate constraining* and 50% of the *inanimate constraining* verbs appeared in discourse contexts that, in conjunction with the verb, constrained strongly for a specific upcoming noun (*high discourse constraint*; mean constraint: 68% +/− 15%), while 50% of the *animate constraining* and 50% of the *inanimate constraining* verbs appeared in discourse contexts that did not constrain strongly for a specific noun (*low discourse constrain*t; mean constraint: 22% +/− 12%). As shown in Figure 3B, the spatial similarity effect appeared to be equally large following the *high discourse constraint* (Figure 3B: left) and the *low discourse constraint* contexts (Figure 3B: right). To statistically quantify this, we averaged the spatial similarity values between 500 - 600ms relative to verb onset (when the effect was maximal) separately for each of the four conditions and used these values as the dependent measure in a repeated measures ANOVA in which Verb animacy constraint (*animate constraining*, *inanimate constrainin*g) and Discourse constraint (*high discourse constraint*, *low discourse constraint*) served as within-subjects factors. This analysis confirmed a main effect of Verb animacy constraint (F_(1,31)_ = 12.05, *p* = .002, η^2^ = 0.28), but failed to reveal either an interaction between Verb animacy constraint and Discourse constraint (F(1,31) = 0.20, *p* = .66, η^2^ = 0.01), or a main effect of Discourse constraint (F_(1,31)_ = 2.43, *p* = .13, η^2^ = 0.07).

#### Spatial similarity results of the EEG data

Figure 5A (left) presents the group-averaged EEG time series of spatial similarity values following *animate constraining* and *inanimate constraining* verbs (averaged across participants from both EEG datasets: 72 in total). Similar to MEG, the overall spatial similarity appeared to increase rapidly from ∼50ms after verb onset, with two sharp peaks at ∼100ms and ∼200ms post verb onset, and then a relatively lower and broader peak between 300 - 400ms following verb onset. Following the onset of the determiner, we observed a similar rapid increase, with three sharp peaks at ∼50ms, ∼175ms and ∼200ms post determiner onset, but no obvious peak between 300 - 400ms. Once again, these overall increases in similarity values appeared to mirror the evoked responses elicited by the verbs and the following determiners, shown at two representative electrode sites in Figure 6.

**Figure 5.**
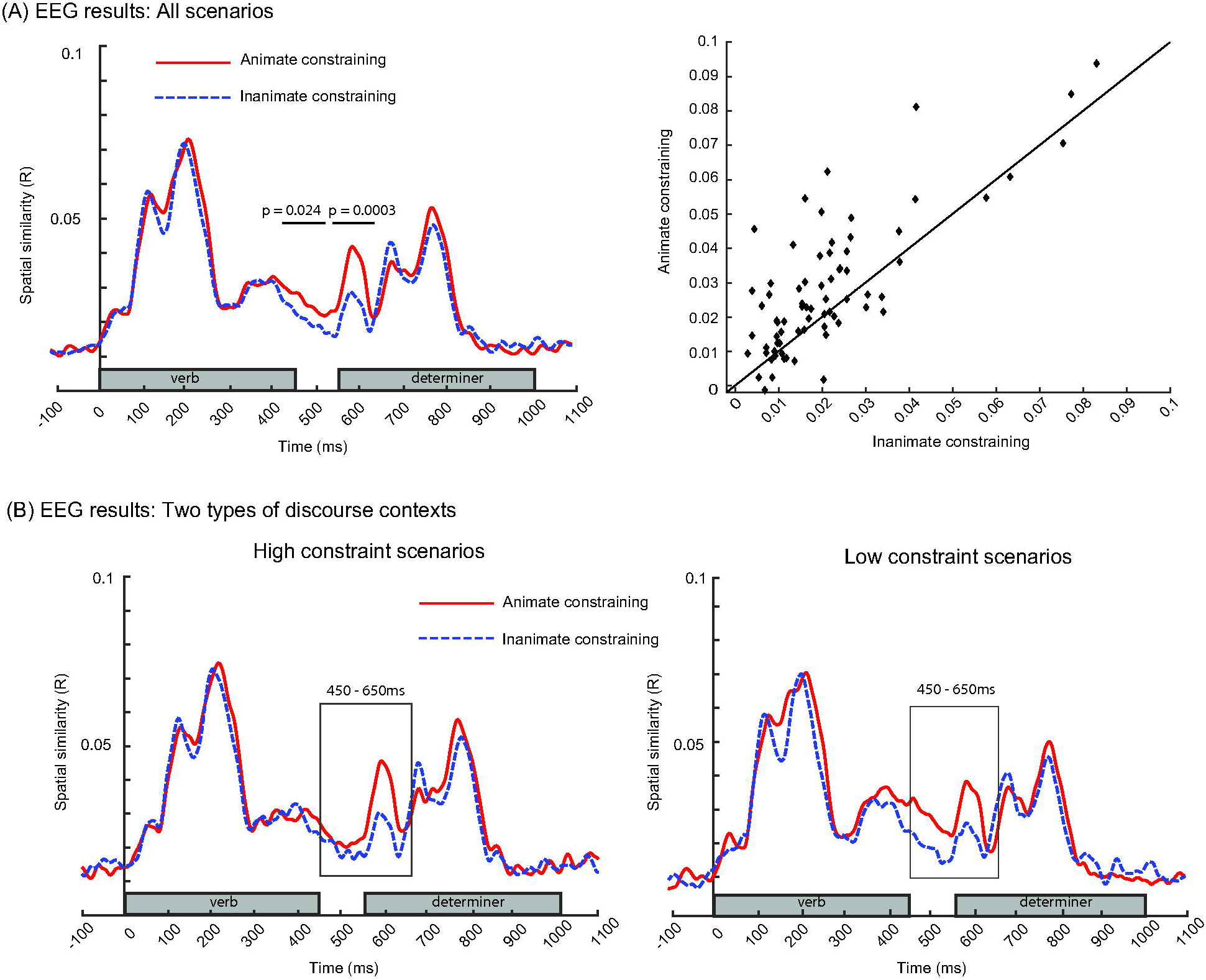
Results of the spatial similarity analysis of the EEG data (combined across the two EEG datasets, 72 participants). (A) Left: Group-averaged time series of spatial similarity values following *animate constraining* verbs (red solid line) and following *inanimate constraining* verbs (blue dotted line), from verb onset at 0ms to noun onset at 1100ms. The duration of the verbs (0 - 450ms) and the subsequent determiners (550 - 1000ms) are marked with grey bars on the x-axis. The spatial pattern of neural activity was more similar following the *animate constraining* than the *inanimate constraining* verbs between 420 - 512ms (*p* = .024) and between 530 - 636ms (*p* = .0003) following verb onset (10000 times permutations); the significant cluster is highlighted by a black line over the time series. Right: A scatter plot of the averaged R-values per participant across the 500 - 600ms time window following verb onset. This shows that two thirds of participants had R-values above the diagonal line, i.e. larger spatial similarity values following the *animate constraining* than the *inanimate constraining* verbs. (B) Group-averaged time series of spatial similarity values following *animate constraining* (red solid line) and *inanimate constraining* (blue dotted line) in the *high discourse constraint* scenarios that constrained strongly for a specific word (left) and the *low discourse constraint* scenarios that did not constrain strongly for a specific noun (right). The spatial similarity effect was equally large following the two types of *discourse constraint* contexts, as indicated by the absence of an interaction between Verb animacy constraint and Discourse constraint (F(1,71) = 0.42, *p* = .52, η^2^ = 0.01) for the averaged spatial similarity values between 450 - 650ms following verb onset.

**Figure 6.**
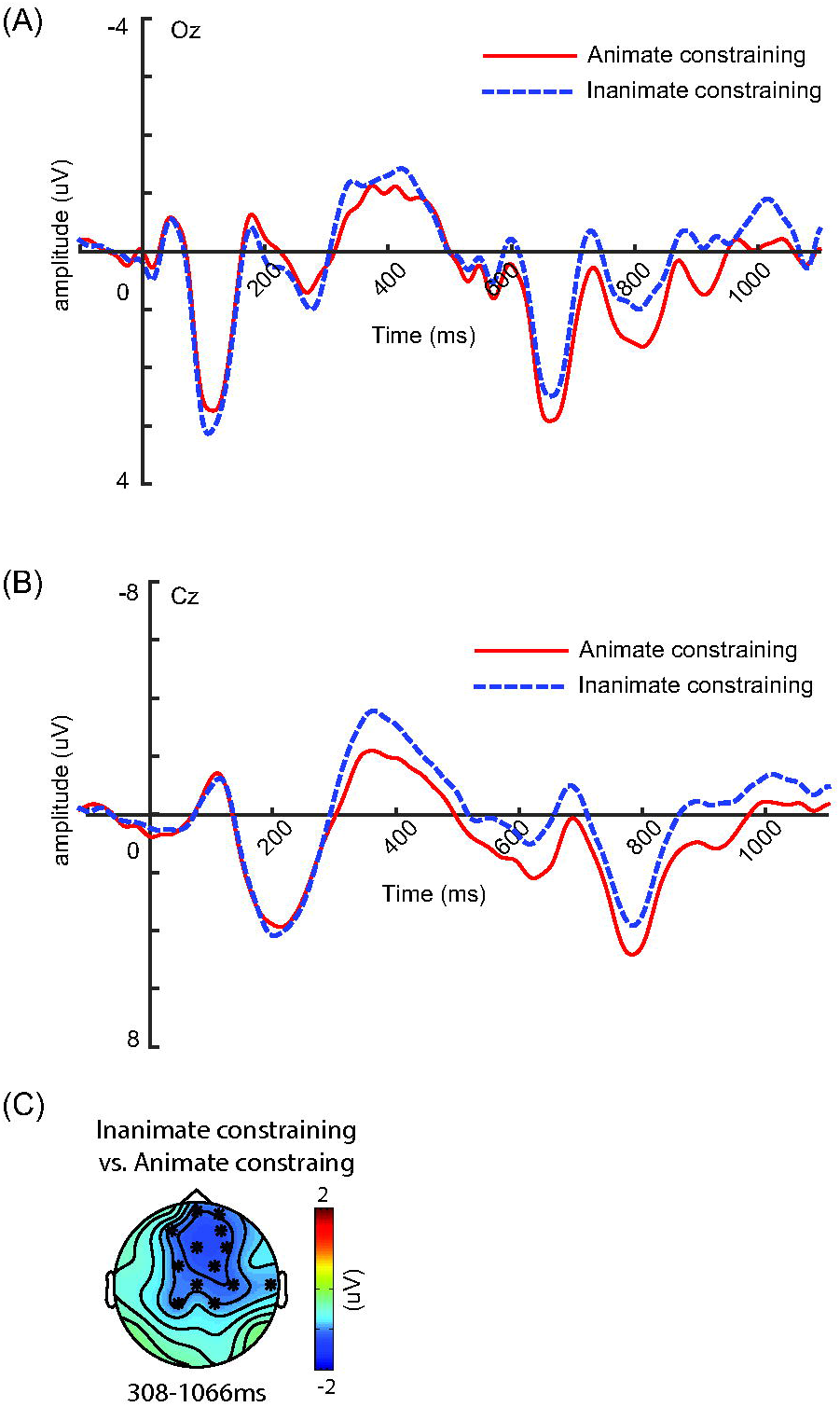
Grand-average event-related potentials (ERPs) of the EEG data (n=72), time-locked to verb onset. ERPs following the onset of *animate constraining* and *inanimate constraining* verbs are shown at two representative electrodes — (A) a midline posterior electrode Oz, and (B) a midline central electrode Cz. As shown in (A), there were strong stimulus-evoked responses that peaked at ∼50ms and ∼100ms following the onset of both the verb and the determiner (at ∼600ms and ∼650ms following verb onset) — the C1 and P1 components that are classically evoked by visual inputs. As shown in (B), there were strong evoked responses that peaked at 100ms (the N1 component), at 200ms (the P2 component) following both the verb and determiner (i.e. at 650ms and 750ms relative to verb onset). Following the verb but not the determiner, the N400 component was observed, peaking at 400ms. (C) The topographic distribution of the ERP difference within the time window where a significant difference was found (*p* = .002, between 308 - 1066ms) based on a cluster-based permutation test over the entire time window. The ERPs evoked by the *inanimate constraining* verbs were larger (more negative) than that evoked by the *animate constraining* verbs at frontal-central EEG electrodes. The electrodes that showed significant differences between the two conditions within the 308 - 1066ms time window are indicated with black asterisks on the topographic map.

Again, of most theoretical interest was whether the spatial similarity pattern of neural activity differed following the *animate constraining* versus the *inanimate constraining* verbs. As shown in Figure 5 (left), similar to MEG, there did indeed appear to be a difference, with larger spatial similarity values following the *animate constraining* than following the *inanimate constraining* verbs from ∼400ms after verb onset. This effect again continued into the interstimulus interval, lasting until around 100ms after determiner onset. A cluster-based permutation test (Maris and Oostenveld, 2007) across the entire epoch (from 0 to 1100ms relative to the onset of verbs) confirmed this difference, revealing two significant clusters between 420 - 512ms, *p* = .024, and between 530 - 636ms, *p* = .0003 relative to the verb onset. Two thirds of participants showed greater spatial similarity values following the *animate constraining* than the *inanimate constraining* verbs within the 450 - 650ms time window (see Figure 5A, right).

Once again, we compared the ERPs of the two conditions to determine whether the spatial similarity effect could be explained by the evoked responses to the verbs (Figure 6). Although there was an ERP difference between the *animate constraining* and *inanimate constraining* verbs (*p* = .002, with a cluster between 308 - 1066ms), this effect had a different time course and went in the opposite direction to the spatial similarity effect: *inanimate constraining* verbs evoked a larger (more negative) response than the *animate constraining* verbs at frontal-central EEG channels — an effect that was likely driven by the greater concreteness of the *inanimate constraining* verbs (mean: 3.33; SD: 0.72; based on Brysbaert et al., 2014) than the *animate constraining* verbs (mean: 2.67; SD: 0.73), t_(173)_ = 5.98, *p* < .001), see Holcomb et al., (1999) and Barber et al., (2013). Importantly, the *similarity structure* of the concreteness values (as tested on the item pairwise difference values) was matched between the two groups of verbs (*p* = .94).

Just as for the MEG dataset, we also asked whether the spatial similarity effect was modulated by the lexical constraint of the broader discourse context. We calculated the spatial similarity time series separately for the *animate constraining* and *inanimate constraining* verbs in the *high discourse constraint* and the *low discourse constraint* contexts (see Figure 5B). Then, in each condition, we averaged the spatial similarity values between 450-650ms (where the spatial similarity effect was maximal), and entered the averaged values into a repeated-measures ANOVA. Again, this analysis confirmed the main effect of Verb animacy constraint (F_(1,71)_ = 23.65, *p* < .001, η^2^ = 0.25), but showed no interaction between Verb animacy constraint and Discourse constraint (F_(1,71)_ = 0.42, *p* = .52, η^2^ = 0.01), and no main effect of Discourse constraint (F_(1,71)_ = 0.22, *p* = .64, η^2^ = 0.003).

We also asked whether the observed spatial similarity effect differed between the two EEG datasets by carrying out an additional ANOVA with spatial similarity values averaged between 450 - 650ms as the dependent measure. In this analysis, Dataset (dataset 1, dataset 2) was a between-subject factor, while Verb animacy constraint (*animate constraining*, *inanimate constrainin*g) and Discourse constraint (*high discourse constraint, low discourse constraint*) were within-subjects factors. This analysis revealed a significant main effect of Verb animacy constraint (F_(1,70)_ = 22.28, *p* < .001, η^2^ = 0.24) as well as a significant interaction between Dataset and Verb animacy constraint (F_(1,70)_ = 5.15, *p* = .026, η^2^ = 0.07). Follow-up analyses in each dataset separately showed a near-significant main effect of Verb animacy constraint in the first dataset, F_(1,31)_ = 3.58, *p* = .068, η^2^ = 0.10, and a more robust main effect of Verb animacy in the second dataset, F_(1,39)_ = 22.99, *p* < .001, η^2^ = 0.37. No other interactions were found.

#### Summary

In both the MEG and the EEG datasets, the timing of the *overall* spatial similarity values (regardless of condition) appeared to broadly mirror the timing of the evoked responses produced by the verb and the following determiner. This is not surprising. As stimulus-evoked activity flows across the cortex, it activates different regions at different latencies, producing a dynamically changing magnetic or electrical field that is detected by MEG or EEG channels at the scalp surface. A large stimulus-induced ERF/ERP will be observed at a given latency because, across multiple trials, stimulus-induced neural activity at this latency will be more consistent in both its phase and in the group of channels to which it projects, compared to at rest. Spatial similarity values are likely to be largest when ERF/ERP components are largest because, across trials, the same underlying stimulus-induced activity will result in a particular spatial pattern of activity (detected across all channels) that will be more similar to each other than at rest when no evoked activity is present. For example, a large P1 component will reflect the fact that, across many light flashes, at 100ms, activity from the occipital cortex will be consistent in its phase and in the subset of channels to which it projects maximally, and this will coincide with a large overall spatial similarity value at 100ms because the overall spatial pattern of activity (more activity at posterior than anterior channels) produced by each flash of light will be more similar to one another than the spatial patterns observed at rest.

Of most theoretical interest was the greater similarity amongst spatial patterns of neural activity following the *animate constraining* than following the *inanimate constraining* verbs in both the MEG and EEG datasets. In both datasets, this effect was significant between 500 - 600ms after verb onset. It cannot be explained by differences in ERFs/ERPs across conditions, and it began *after* the peak of N400 component evoked by the verb, and *after* the overall spatial similarity values had begun to decrease. These observations support our interpretation that it reflected *anticipatory* activity for the upcoming noun that was not directly linked to bottom-up activity evoked by the verb itself.

The strikingly convergent findings across the EEG and MEG RSA analyses are consistent with a recent study that used RSA together with both EEG and MEG to decode visual representations of living versus non-living objects (Cichy and Pantazis, 2017). MEG and EEG are sensitive to neural activity from different underlying sources (e.g. MEG is most sensitive to activity originating within sulci, while EEG is sensitive to activity originating in both sulci and gyri). However, both methods are able to detect post-synaptic activity produced by pyramidal cells within highly distributed cortical locations, giving rise to spatial patterns of activity on the surface of the scalp. Our findings suggest that, with both techniques, RSA was able to capture differences between our experimental conditions in the similarity amongst these spatial patterns. It is particularly encouraging that, just as in the study described by Cichy and Pantazis (2017), we showed that EEG RSA was able to discriminate between the animate and inanimate conditions, despite the fact that the EEG signal is more smeared at the surface of the scalp than MEG (Hämäläinen et al., 1993), and that we used fewer channels to collect our EEG data (in the first MEG-EEG study: 70 electrodes; in the second EEG-only study: 32 electrodes) than our MEG data in the first study (306 sensors).

### Behavioral findings

We did not acquire behavioral data on the verb itself. However, in both experiments, at the end of each scenario, participants made acceptability judgments, with acceptability determined by whether the direct object noun matched or violated the animacy constraints of the verb. In the MEG-EEG study, participants made correct judgments in 84.09% of scenarios on average (SD: 7.32%), with no differences between scenarios that contained *animate constraining* and *inanimate constraining* verbs (t_(31)_ = 1.60, *p* = .12). In the EEG-only study, participants made correct judgments in 89.17% of scenarios on average (SD: 5.26%), again with no differences between scenarios containing *animate constraining* and *inanimate constraining* verbs (t_(31)_ = 0.71, *p* = .48).

In addition to making acceptability judgments after each scenario, participants also responded to Yes/No questions that followed a subset of scenarios. In the MEG-EEG study, on average, 76.56% of the 24 comprehension questions were answered correctly (SD: 16.18%), and in the EEG-only study, 84.94% of the 32 comprehension questions were answered correctly (SD: 6.75%). These findings indicate that participants attended to the context information within the discourse scenarios, rather than only to the final sentences.

## Discussion

We conducted a spatial similarity analysis on MEG and EEG data to ask whether comprehenders use the semantic constraints of verbs to predict the animacy of upcoming nouns during sentence comprehension. Our findings were robust and strikingly convergent across the MEG (n=32) and EEG (n=72) datasets. The spatial pattern of neural activity following *animate constraining* verbs was significantly more similar than following *inanimate constraining* verbs. This effect started to emerge at around 450ms (EEG)/500ms (MEG) following verb onset — past the peak of the evoked N400 produced by the verb. It is therefore unlikely to have reflected differences in lexico-semantic processing of the verb itself. It also cannot be explained by differences between the *animate constraining* and *inanimate constraining* verbs in aspects of their semantic and/or lexical similarity structures that were unrelated to their following arguments, as these were matched across conditions (see Methods).

In general, verbs that constrain for animate direct object nouns also constrain for fewer types of thematic/syntactic structures than verbs that constrain for inanimate nouns (Kipper et al., 2006). Therefore, in theory, the differences in spatial similarity following the *animate constraining* versus *inanimate constraining* verbs could have reflected differences in the *syntactic* similarity structure of the predicted upcoming inputs. In this study, however, all the verbs had a transitive bias, and they all appeared in the same subject-verb-noun syntactic structure. We therefore think that, just as in the second cloze-norming study, comprehenders predicted direct object nouns in all sentences, and that the spatial similarity effect was driven by differences in the *semantic* similarity structure of the predicted *animate* and *inanimate* upcoming nouns.

There has been much debate about how we are able to make categorical distinctions based on animacy. One set of proposals assumes that *animate* and *inanimate* concepts are encoded within distinct neural systems that are separated based on either categorical domain (animacy, e.g. Caramazza and Shelton, 1998) or modality (perceptual features for animate concepts and functional properties for inanimate concepts; e.g. Warrington and Shallice, 1984). These accounts are supported by functional neuroimaging and EEG studies reporting spatially distinct patterns of neural activity in response to animate versus inanimate stimuli (e.g. Martin et al., 1996; Sitnikova et al., 2006). However, the neuroanatomical location of this activity tends to be quite inconsistent across studies (Tyler and Moss, 2001). Moreover, these types of ‘localization’ accounts cannot explain how animacy-based categorization deficits arise in patients with non-focal neuropathologies such as Alzheimer’s disease (e.g. Gonnerman et al., 1997).

An alternative explanation is that, instead of reflecting distinct localizable stores of knowledge, the animate-inanimate distinction emerges implicitly from differences in the degree of similarity amongst the sets of *distributed* semantic features/attributes that characterize animate and inanimate concepts, which are represented across widespread regions of the cortex (Devlin et al., 1998; Taylor et al., 2011). Highly distributed patterns of cortical activity give rise to distinct spatial patterns of electrical and magnetic activity detected by EEG/MEG channels at the surface of the scalp. The greater intercorrelation amongst the semantic properties that characterize animate concepts than inanimate concepts will therefore be reflected by a greater intercorrelation amongst the spatial patterns of EEG/MEG activity associated with their processing. These differences can be detected using RSA (Cichy et al., 2014; Stokes et al., 2015), and it is precisely this sensitivity to spatial similarity structure that, in previous studies, allowed animacy to be decoded from spatial patterns of EEG/MEG activity produced by bottom-up linguistic (Sudre et al., 2012) and non-linguistic (Carlson et al., 2013; Cichy et al., 2014; Cichy and Pantazis, 2017; Khaligh-Razavi et al., 2018) inputs. Here, we show for the first time that RSA can be used in combination with EEG/MEG to detect distinct spatial similarity patterns during language comprehension *before* new bottom-up inputs become available, reflecting the *pre-activation* of animacy-linked semantic features.

### The prediction of upcoming animacy features was not dependent on the prediction of a specific word

While in previous work, we have combined MEG and RSA to show anticipatory neural activity associated with the prediction of specific upcoming individual words (Wang et al., 2018), the present findings provide neural evidence for the pre-activation of semantic features that characterize whole sets of words. We further showed that predicting these broad sets of semantic features did not depend on being able to predict a single word: the spatial similarly effect was just as large following low constraint as following high constraint discourse contexts.

This finding has important theoretical implications. It has sometimes been argued that, because most words are not highly predictable on the basis of their prior contexts, predictive processing is unlikely to play a major role in language comprehension. Implicit in this argument is the assumption that we are only able to predict upcoming lexical items. We and others, however, have argued that comprehenders are able to predict upcoming information, with various degrees of certainty, at multiple levels and grains of representation (e.g. Altmann and Mirković, 2009; Kuperberg and Jaeger, 2016). The present findings show that, despite not being able to predict upcoming words, the constraints of the verb provided enough information for comprehenders to predict upcoming semantic features that distinguished between upcoming *animate* and *inanimate* items (see also Szewczyk and Schriefers, 2013). More generally, by showing that the combination of RSA with EEG/MEG can detect pre-activated semantic representations in the absence of new bottom-up inputs, our findings suggest that this combination can be used to examine whether we predict finer-grained semantic categories during language comprehension. For example, following the contexts, “Her favorite vegetable is …” and “The carpenter is making a …”, it will be interesting to determine whether we pre-activate distinct patterns of neural activity that correspond to the predicted <vegetables> and <furniture> categories respectively — categories that are known to have distinct semantic similarity structures (Cree and McRae, 2003), which can be decoded from brain activity (Kriegeskorte et al., 2008b).

### The time course of the prediction effect

As noted above, the spatial similarity effect began past the stage at which comprehenders are likely to have accessed other lexico-semantic features of the verb, and well before the argument actually appeared. We suggest that this was the first time point at which comprehenders were able to infer the full high-level event structure (e.g. <Agent cautioned animate noun>), and that they used this structure to generate top-down predictions of the semantic features linked to the animacy of upcoming arguments (Kuperberg and Jaeger, 2016; Kuperberg et al., 2019).

Despite its early onset, the spatial similarity effect lasted for only around 150ms (MEG)/200ms (EEG). This is consistent with a recent MEG-RSA study in which we used a different paradigm in a different language (Chinese) to capture the prediction of specific individual words (Wang et al., 2018). These types of short-lived prediction effects might seem surprising if one assumes that pre-activated mental representations are necessarily accompanied by sustained detectable neural activity. However, evidence from intracranial recordings of local neural activity (Mongillo et al., 2008; Stokes et al., 2013; Lundqvist et al., 2016; Bastos et al., 2018; Lundqvist et al., 2018b), and from noninvasive EEG and fMRI recordings of global brain activity (Sprague et al., 2016; Wolff et al., 2017), suggests that, instead of being persistent, neural activity over delays can be relatively sparse, especially when other information is concurrently activated from long term memory (Kaminski et al., 2017). During these delays, anticipated information remains accessible, but it can only be *detected* when perturbed or “pinged”, e.g. by a targeted pulse of transcranial magnetic stimulation (Rose et al., 2016), or by new bottom-up input (Wolff et al., 2017). This has led to the hypothesis that anticipated information is held in an “activity silent” state (Stokes, 2015; Lundqvist et al., 2018a), becoming available only when it is task relevant (Sprague et al., 2016; Lundqvist et al., 2018b).

Extrapolating to the present findings, we speculate that, despite the absence of a spatial similarity effect immediately preceding the noun, the predicted animacy-linked semantic features were nonetheless available to facilitate semantic processing of the incoming noun when it appeared. And, indeed, as in many previous studies (e.g. Paczynski and Kuperberg, 2011, 2012; Szewczyk and Schriefers, 2011; Kuperberg, et al., 2019), the evoked N400 response on the subsequent noun was reduced when its animacy features matched (versus mismatched) the animacy constraints of the preceding verb. Moreover, a spatial similarity analysis on the subsequent noun confirmed that the spatial pattern of neural activity was more similar to plausible animate than inanimate nouns (Wang and Kuperberg, Unpublished).

In sum, we provide direct neural evidence for the prediction of animacy-linked semantic features during the comprehension of short discourse scenarios. These findings pave the way towards combining RSA with EEG/MEG to yield insights into the nature and specificity of prediction, and its neural instantiation, during language comprehension.

## Acknowledgements

This work was funded by the National Institute of Child Health and Human Development (R01 HD08252 to G.R.K.). We thank Nao Matsuda, Sheraz Khan and Matt Hämäläinen for technical MEG support. We thank Maria Luiza, Cunha Lima, Margarita Zeitlin, and Connie Choi for their contributions to constructing the experimental materials, Margarita Zeitlin, Simone Riley, Eric Fields, Allison Fogel and Arim Choi for their assistance with data collection, Rebeca Becdach for her assistance with identifying senses of stimuli, as well as Ole Jensen and Trevor Brothers for helpful discussions.

## References

Altmann GT, Kamide Y (1999) Incremental interpretation at verbs: Restricting the domain of subsequent reference. Cognition 73:247–264.

Altmann GT, Mirković J (2009) Incrementality and prediction in human sentence processing. Cognitive Science 33:583–609.

Balota DA, Yap MJ, Hutchison KA, Cortese MJ, Kessler B, Loftis B, Neely JH, Nelson DL, Simpson GB, Treiman R (2007) The English lexicon project. Behavior Research Methods 39:445–459.

Barber HA, Otten LJ, Kousta S-T, Vigliocco G (2013) Concreteness in word processing: ERP and behavioral effects in a lexical decision task. Brain and language 125:47–53.

Bastos AM, Loonis R, Kornblith S, Lundqvist M, Miller EK (2018) Laminar recordings in frontal cortex suggest distinct layers for maintenance and control of working memory. Proceedings of the National Academy of Sciences 115:1117–1122.

Bell AJ, Sejnowski TJ (1997) The “independent components” of natural scenes are edge filters. Vision Research 37:3327–3338.

Brysbaert M, New B (2009) Moving beyond Kućera and Francis: A critical evaluation of current word frequency norms and the introduction of a new and improved word frequency measure for American English. Behavior Research Methods 41:977–990.

Brysbaert M, Warriner AB, Kuperman V (2014) Concreteness ratings for 40 thousand generally known English word lemmas. Behavior Research Methods 46:904–911.

Caramazza A, Shelton JR (1998) Domain-specific knowledge systems in the brain: The animate-inanimate distinction. Journal of Cognitive Neuroscience 10:1–34.

Carlson T, Tovar DA, Alink A, Kriegeskorte N (2013) Representational dynamics of object vision: the first 1000 ms. Journal of Vision 13:1–1.

Cichy RM, Pantazis D (2017) Multivariate pattern analysis of MEG and EEG: A comparison of representational structure in time and space. NeuroImage 158:441–454.

Cichy RM, Pantazis D, Oliva A (2014) Resolving human object recognition in space and time. Nature Neuroscience 17:455–462.

Cree, GS, & McRae, K (2003). Analyzing the factors underlying the structure and computation of the meaning of chipmunk, cherry, chisel, cheese, and cello (and many other such concrete nouns). Journal of Experimental Psychology: General, 132(2), 163.

Dahl Ö (2008) Animacy and egophoricity: Grammar, ontology and phylogeny. Lingua 118:141–150.

Devlin JT, Gonnerman LM, Andersen ES, Seidenberg MS (1998) Category-specific semantic deficits in focal and widespread brain damage: A computational account. Journal of Cognitive Neuroscience 10:77–94.

Federmeier KD, Wlotko EW, De Ochoa-Dewald E, Kutas M (2007) Multiple effects of sentential constraint on word processing. Brain Research 1146:75–84.

Fellbaum C (1990) English Verbs as a Semantic Net. International Journal of Lexicography 3:278–301.

Ferretti TR, McRae K, Hatherell A (2001) Integrating verbs, situation schemas, and thematic role concepts. Journal of Memory and Language 44:516–547.

Garrard P, Lambon Ralph MA, Hodges JR, Patterson K (2001) Prototypicality, distinctiveness, and intercorrelation: Analyses of the semantic attributes of living and nonliving concepts. Cognitive Neuropsychology 18:125–174.

Gonnerman LM, Andersen ES, Devlin JT, Kempler D, Seidenberg MS (1997) Double dissociation of semantic categories in Alzheimer’s disease. Brain and language 57:254–279.

Gramfort A, Luessi M, Larson E, Engemann DA, Strohmeier D, Brodbeck C, Parkkonen L, Hämäläinen MS (2014) MNE software for processing MEG and EEG data. NeuroImage 86:446–460.

Gramfort A, Luessi M, Larson E, Engemann D, Strohmeier D, Brodbeck C, Goj R, Jas M, Brooks T, Parkkonen L, Hämäläinen M (2013) MEG and EEG data analysis with MNE-Python. Frontiers in Neuroscience 7.

Hämäläinen M, Hari R, Ilmoniemi RJ, Knuutila J, Lounasmaa OV (1993) Magnetoencephalography—theory, instrumentation, and applications to noninvasive studies of the working human brain. Reviews of Modern Physics 65:413.

Hare M, Jones M, Thomson C, Kelly S, McRae K (2009) Activating event knowledge. Cognition 111:151–167.

Holcomb PJ, Kounios J, Anderson JE, West WC (1999) Dual-coding, context-availability, and concreteness effects in sentence comprehension: An electrophysiological investigation. Journal of Experimental Psychology: Learning, Memory, and Cognition 25:721.

Huth AG, De Heer WA, Griffiths TL, Theunissen FE, Gallant JL (2016) Natural speech reveals the semantic maps that tile human cerebral cortex. Nature 532:453.

Jackendoff R (1993) On the role of conceptual structure in argument selection: A reply to Emonds. Natural Language & Linguistic Theory 11:279–312.

Jung T-P, Makeig S, Westerfield M, Townsend J, Courchesne E, Sejnowski TJ (2000) Removal of eye activity artifacts from visual event-related potentials in normal and clinical subjects. Clinical Neurophysiology 111:1745–1758.

Kaminski J, Sullivan S, Chung JM, Ross IB, Mamelak AN, Rutishauser U (2017) Persistently active neurons in human medial frontal and medial temporal lobe support working memory. Nature Neuroscience 20:590–601.

Kemp C, Tenenbaum JB (2008) The discovery of structural form. Proceedings of the National Academy of Sciences 105:10687–10692.

Khaligh-Razavi S-M, Cichy RM, Pantazis D, Oliva A (2018) Tracking the spatiotemporal neural dynamics of real-world object size and animacy in the human brain. Journal of Cognitive Neuroscience 30:1559–1576.

Kipper K, Korhonen A, Ryant N, Palmer M (2006) Extending VerbNet with novel verb classes. In: Proceedings of LREC, p 1: Citeseer.

Kriegeskorte N, Mur M, Bandettini P (2008a) Representational similarity analysis – connecting the branches of systems neuroscience. Frontiers in Systems Neuroscience 2:4.

Kriegeskorte N, Mur M, Ruff DA, Kiani R, Bodurka J, Esteky H, Tanaka K, Bandettini PA (2008b) Matching categorical object representations in inferior temporal cortex of man and monkey. Neuron 60:1126–1141.

Kuperberg GR, Jaeger TF (2016) What do we mean by prediction in language comprehension? Language, Cognition and Neuroscience 31:32–59.

Kuperberg GR, Brothers T, Wlotko EW (2019) A tale of two positivities and the N400: Distinct neural signatures are evoked by confirmed and violated predictions at different levels of representation. Journal of Cognitive Neuroscience 32:12–35.

Levin B (1993) English verb classes and alternations: A preliminary investigation: University of Chicago press.

Loper E, Bird S (2002) NLTK: the Natural Language Toolkit. In: Proceedings of the ACL-02 Workshop on effective tools and methodologies for teaching natural language processing and computational linguistics - Volume 1, pp 63–70. Philadelphia, Pennsylvania: Association for Computational Linguistics.

Luke SG, Christianson K (2016) Limits on lexical prediction during reading. Cognitive Psychology 88:22–60.

Lund K, Burgess C (1996) Producing high-dimensional semantic spaces from lexical co-occurrence. Behavior Research Methods, Instruments, & Computers 28:203–208.

Lundqvist M, Herman P, Miller EK (2018a) Working memory: delay activity, yes! persistent activity? Maybe not. Journal of Neuroscience 38:7013–7019.

Lundqvist M, Herman P, Warden MR, Brincat SL, Miller EK (2018b) Gamma and beta bursts during working memory readout suggest roles in its volitional control. Nature Communications 9:394.

Lundqvist M, Rose J, Herman P, Brincat Scott L, Buschman Timothy J, Miller Earl K (2016) Gamma and beta bursts underlie working memory. Neuron 90:152–164.

Maris E, Oostenveld R (2007) Nonparametric statistical testing of EEG- and MEG-data. Journal of Neuroscience Methods 164:177–190.

Martin A (2016) GRAPES—Grounding representations in action, perception, and emotion systems: How object properties and categories are represented in the human brain. Psychonomic Bulletin & Review 23:979–990.

Martin A, Wiggs CL, Ungerleider LG, Haxby JV (1996) Neural correlates of category-specific knowledge. Nature 379:649.

McRae K, De Sa VR, Seidenberg MS (1997) On the nature and scope of featural representations of word meaning. Journal of Experimental Psychology: General 126:99.

Miller GA (1990) Nouns in WordNet: a lexical inheritance system. International Journal of Lexicography 3:245–264.

Miller GA, Beckwith R, Fellbaum C, Gross D, Miller KJ (1990) Introduction to WordNet: An on-line lexical database. International Journal of Lexicography 3:235–244.

Mongillo G, Barak O, Tsodyks M (2008) Synaptic theory of working memory. Science 319:1543–1546.

Nairne JS, VanArsdall JE, Cogdill M (2017) Remembering the living: Episodic memory is tuned to animacy. Current Directions in Psychological Science 26:22–27.

Oldfield RC (1971) The assessment and analysis of handedness: the Edinburgh inventory. Neuropsychologia 9:97–113.

Oostenveld R, Fries P, Maris E, Schoffelen J-M (2011) FieldTrip: open source software for advanced analysis of MEG, EEG, and invasive electrophysiological data. Computational Intelligence and Neuroscience 2011:1.

Paczynski M, Kuperberg GR (2011) Electrophysiological evidence for use of the animacy hierarchy, but not thematic role assignment, during verb argument processing. Language and Cognitive Processes 26:1402–1456.

Paczynski M, Kuperberg GR (2012) Multiple influences of semantic memory on sentence processing: Distinct effects of semantic relatedness on violations of real-world event/state knowledge and animacy selection restrictions. Journal of Memory and Language 67:426–448.

Peirce JW (2007) PsychoPy--Psychophysics software in Python. Journal Neuroscience Methods 162:8–13.

Perrin F, Pernier J, Bertrand O, Echallier J (1989) Spherical splines for scalp potential and current density mapping. Electroencephalography and Clinical Neurophysiology 72:184–187.

Randall B, Moss HE, Rodd JM, Greer M, Tyler LK (2004) Distinctiveness and correlation in conceptual structure: Behavioral and computational studies. Journal of Experimental Psychology: Learning, Memory, and Cognition 30:393.

Rogers TT, McClelland JL (2008) Précis of semantic cognition: A parallel distributed processing approach. Behavioral and Brain Sciences 31:689–714.

Rose NS, LaRocque JJ, Riggall AC, Gosseries O, Starrett MJ, Meyering EE, Postle BR (2016) Reactivation of latent working memories with transcranial magnetic stimulation. Science 354:1136–1139.

Schwanenflugel PJ, LaCount KL (1988) Semantic relatedness and the scope of facilitation for upcoming words in sentences. Journal of Experimental Psychology: Learning, Memory, and Cognition 14:344.

Sitnikova T, West WC, Kuperberg GR, Holcomb PJ (2006) The neural organization of semantic memory: Electrophysiological activity suggests feature-based segregation. Biological Psychology 71:326–340.

Sprague TC, Ester EF, Serences JT (2016) Restoring latent visual working memory representations in human cortex. Neuron 91:694–707.

Stokes MG, Kusunoki M, Sigala N, Nili H, Gaffan D, Duncan J (2013) Dynamic coding for cognitive control in prefrontal cortex. Neuron 78:364–375.

Stokes MG, Wolff MJ, Spaak E (2015) Decoding rich spatial information with high temporal resolution. Trends in Cognitive Sciences 19:636–638.

Sudre G, Pomerleau D, Palatucci M, Wehbe L, Fyshe A, Salmelin R, Mitchell T (2012) Tracking neural coding of perceptual and semantic features of concrete nouns. NeuroImage 62:451–463.

Szewczyk JM, Schriefers H (2011) Is animacy special?: ERP correlates of semantic violations and animacy violations in sentence processing. Brain Research 1368:208–221.

Szewczyk JM, Schriefers H (2013) Prediction in language comprehension beyond specific words: An ERP study on sentence comprehension in Polish. Journal of Memory and Language 68:297–314.

Taylor KI, Devereux BJ, Tyler LK (2011) Conceptual structure: Towards an integrated neurocognitive account. Language and Cognitive Processes 26:1368–1401.

Tyler LK, Moss HE (2001) Towards a distributed account of conceptual knowledge. Trends in cognitive sciences 5:244–252.

Taylor WL (1953) “Cloze procedure”: A new tool for measuring readability. Journalism Bulletin 30:415–433.

Tyler LK, Moss HE (2001) Towards a distributed account of conceptual knowledge. Trends in Cognitive Sciences 5:244–252.

Uusitalo MA, Ilmoniemi RJ (1997) Signal-space projection method for separating MEG or EEG into components. Medical and Biological Engineering and Computing 35:135–140.

Wang L, Kuperberg G, Jensen O (2018) Specific lexico-semantic predictions are associated with unique spatial and temporal patterns of neural activity. eLife 7:e39061.

Warrington EK, Shallice T (1984) Category specific semantic impairments. Brain 107:829–853.

White K, Ashton R (1976) Handedness assessment inventory. Neuropsychologia 14:261–264.

Wicha NY, Moreno EM, Kutas M (2004) Anticipating words and their gender: An event-related brain potential study of semantic integration, gender expectancy, and gender agreement in Spanish sentence reading. Journal of Cognitive Neuroscience 16:1272–1288.

Wolff MJ, Jochim J, Akyurek EG, Stokes MG (2017) Dynamic hidden states underlying working-memory-guided behavior. Nature Neuroscience 20:864–871.

Wu Z, Palmer M (1994) Verbs semantics and lexical selection. In: Proceedings of the 32nd annual meeting on Association for Computational Linguistics, pp 133–138: Association for Computational Linguistics.

Zannino GD, Perri R, Pasqualetti P, Caltagirone C, Carlesimo GA (2006) Analysis of the semantic representations of living and nonliving concepts: a normative study. Cognitive Neuropsychology 23:515–540.

